# CCR5 deficiency normalizes TIMP levels, working memory, and gamma oscillation power in APOE4 targeted replacement mice

**DOI:** 10.1101/2022.06.29.498148

**Authors:** Griffin A. Greco, Mitchell Rock, Matthew Amontree, Maria Fe Lanfranco, Holly Korthas, Sung Hyeok Hong, R. Scott Turner, G. William Rebeck, Katherine Conant

## Abstract

The *APOE4* allele increases the risk for Alzheimer’s disease (AD) in a dose-dependent manner and is also associated with cognitive decline in non-demented elderly controls. In mice with targeted gene replacement (TR) of murine *APOE* with human *APOE3* or *APOE4*, the latter show reduced neuronal dendritic complexity and impaired learning. *APOE4* TR mice also show reduced gamma oscillation power and sharp wave ripple (SWR) abundance, neuronal population activities important to learning and memory. Published work has shown that brain extracellular matrix (ECM) can reduce neuroplasticity as well as gamma power and SWR abundance, while attenuation of ECM can instead enhance these endpoints. In the present study we examine human cerebrospinal fluid (CSF) samples from *APOE3* and *APOE4* individuals and brain lysates from *APOE3* and *APOE4* TR mice for levels of ECM effectors that can increase matrix deposition and restrict neuroplasticity. We find that CCL5, a molecule linked to ECM deposition in liver and kidney, is increased in CSF samples from *APOE4* individuals. Levels of tissue inhibitor of metalloproteinases (TIMPs), which inhibit the activity of ECM-degrading enzymes, are also increased in *APOE4* CSF as well as protein lysates from *APOE4* TR mice. Importantly, as compared to wildtype/*APOE4* heterozygotes, CCR5 knockout/*APOE4* heterozygotes show reduced TIMP levels and enhanced EEG gamma power. The latter also show improved learning and memory, suggesting that the CCR5/CCL5 axis could represent a therapeutic target for *APOE4* individuals.

## Introduction

Alzheimer’s disease (AD) is a leading cause of dementia and by 2050, it will affect approximately 50% of individuals by age 85 (Brookmeyer *et al*., 2007). While the most significant risk factor for AD is age, the greatest genetic risk factor for AD is a genetic variant of apolipoprotein E (*APOE*), which encodes a protein responsible for lipid transport (Corder *et al*., 1993; Poirier *et al*., 1993; Rebeck *et al*., 1993; Strittmatter *et al*., 1993).

In contrast to mice, humans have three different isoforms of ApoE: E2, E3, and E4. These differ by a single amino acid substitution at the 112^th^ and 158^th^ positions. The *APOε4* allele enhances the risk for developing AD (Rebeck *et al*., 1993; Strittmatter *et al*., 1993) while the *APOε2* allele diminishes this risk (Poirier *et al*., 1993). Interestingly, *APOE* variants also influence the age of onset of AD, so that *APOE4* carriers tend to get AD at earlier ages as compared to non-E4 carriers (Poirier *et al*., 1993). Consistent with this observation, *APOE* genotype is thought to affect early processes in AD pathogenesis, such as Aβ accumulation or clearance (Kim *et al*., 2009).

*APOE* may also affect brain structure and function independent of amyloid deposition. *APOE4* has been associated with increased cognitive decline in elderly *APOE4* carriers who do not have AD (Plassman *et al*., 1997). Carriers in their 50s-60s, who are cognitively normal, show reduced glucose metabolism in parietal, temporal and prefrontal cortices (Small *et al*., 1995; Caselli *et al*., 2001). *APOE* genotype also alters the volume of the entorhinal cortex prior to the development of AD (Shaw *et al*., 2007). In murine studies, *APOE4* TR mice show neuronal simplification in the amygdala (Wang *et al*., 2005), and cortex (Dumanis *et al*., 2009). Furthermore, *APOE4* TR mice have abnormalities in hippocampal long-term potentiation (LTP) (Trommer *et al*., 2004; Trommer *et al*., 2005; Korwek *et al*., 2009). Importantly, *APOE4* TR mice have reduced SWR abundance and low gamma power, brain rhythms important to memory consolidation and working memory (Gillespie *et al*., 2016; Jones *et al*., 2019). Intriguingly, reduced SWR abundance in these animals occurred at a relatively young age and the magnitude of reduction correlated with the magnitude of later cognitive impairment (Jones *et al*., 2019).

Accumulating evidence suggests that *APOE4* may be associated with low grade inflammation and glial activation (Tai *et al*., 2015). Of interest, both activated astrocytes and microglia secrete CCL5 (Lanfranco *et al*., 2017), a chemotactic cytokine that potently restricts neuroplasticity and LTP (Zhou *et al*., 2016; Shen *et al*., 2022). CCL5 also inhibits pyramidal cell excitability which could in turn modulate gamma power and SWR abundance (Zhou *et al*., 2016).

CCL5 has also been linked to excess ECM deposition (Passman *et al*., 2021; Bonnard *et al*., 2022). Brain ECM exists in diffuse and condensed forms and both types can restrict neuroplasticity (Miyata & Kitagawa, 2017). For example, dendritic arbor is increased in mice with reductions in ECM-dense perineuronal nets (PNNs) (Alaiyed *et al*., 2020), which are predominantly localized to parvalbumin (PV) expressing inhibitory interneurons and increase their excitability through varied mechanisms (Bozzelli *et al*., 2018). Importantly, enzymatic digestion of brain ECM increases the power of *in vivo* cortical gamma oscillations (Lensjo *et al*., 2017) and the abundance of *ex vivo* hippocampal SWRs (Sun *et al*., 2018).

ECM levels are modulated by deposition as well as degradation, and the latter is mediated by metalloproteinases including matrix metalloproteases (MMPs) and <u>A D</u>isintegrin <u>A</u>nd <u>M</u>etalloproteases (ADAMs) (Rivera, 2019). These enzymes are zinc-dependent secreted or transmembrane endoproteases that have been well-studied for their ability to process proteins of the extracellular matrix but are now appreciated to act on a variety of soluble molecules and cell surface receptors as well (Conant *et al*., 2015). Family members that have been particularly well-implicated in neuroplasticity include MMP-2, MMP-3 and MMP-9 (Nagy *et al*., 2006; Okulski *et al*., 2007; Conant *et al*., 2010; Smith *et al*., 2014; Wojtowicz & Mozrzymas, 2014; Murase *et al*., 2017; Wiera *et al*., 2017). MMP and ADAM activity is in turn inhibited by TIMPs.

MMP-9 is of particular interest in that it is neuronal-derived and its expression/release and/or activation is increased with pyramidal neuron activity (Szklarczyk *et al*., 2002). It is released from pre- and post-synaptic stores and thus it likely targets synaptically localized adhesion molecules and glutamate inputs to PNN-enveloped PV cells. MMP-9 activity has been well-implicated in hippocampal dependent learning and memory as well as striatal- and amygdala-based learning (Nagy *et al*., 2006; Ganguly *et al*., 2013; Smith *et al*., 2014). And while MMP-9 levels may be elevated with late-stage AD and correlated with inflammation (Weekman & Wilcock, 2016), late-stage increases are likely due to microglial activation and other events that instead lead to maladaptive enzyme localization at regions including the blood brain barrier.

In terms of the mechanisms by which MMPs can influence learning and memory, ECM and/or PNN independent effects have also been well-described. For example, MMPs can activate pro-neurotrophins (Lee *et al*., 2001). These enzymes can also cleave cell adhesion molecules to generate N-terminal fragments which serve as integrin-binding ligands (Lonskaya *et al*., 2013). Of interest, integrin signaling may be critical to MMP induced LTP and dendritic spine expansion (Wang *et al*., 2008). MMP dependent activation protease activated receptor-1 (PAR-1) can also increase dendritic arborization (Allen *et al*., 2016) as well as some forms of LTP (Almonte *et al*., 2013).

In the present study, our aim was to explore the possibility that CCL5 and other potential ECM effectors that limit pyramidal arbor, gamma power and SWR abundance may be elevated in *APOE4* human and TR mice. We also investigated the possibility that reductions in CCR5, a molecule linked to excess ECM deposition in other end organs as well as reduced plasticity in the brain (Passman *et al*., 2021; Bonnard *et al*., 2022; Shen *et al*., 2022), might rescue biochemical, neurophysiological and/or behavioral alterations associated with this allele.

## Materials and Methods

### Human CSF samples

Human CSF samples were collected for a randomized placebo-controlled double-blind, multi-site, phase 2 trial of resveratrol in individuals with mild to moderate dementia due to AD (Turner *et al*., 2015). Concomitant use of FDA-approved medications for AD (e.g., cholinesterase inhibitors) was allowed. The two randomized groups were similar at baseline with the exception that duration of diagnosis was longer in the placebo group. Participants (total *N* = 119) were randomized to placebo or resveratrol 500 mg orally once daily (with a dose escalation by 500-mg increments every 13 weeks, ending with 1000 mg twice daily). The total treatment duration was 52 weeks. For data presented herein we analyzed the pretreatment samples (Figure 1 a-d), or alternatively after pretreatment stocks were depleted, the 52-week post placebo-treated individuals (Figure 1e and figure 2).

**Figure 1.**
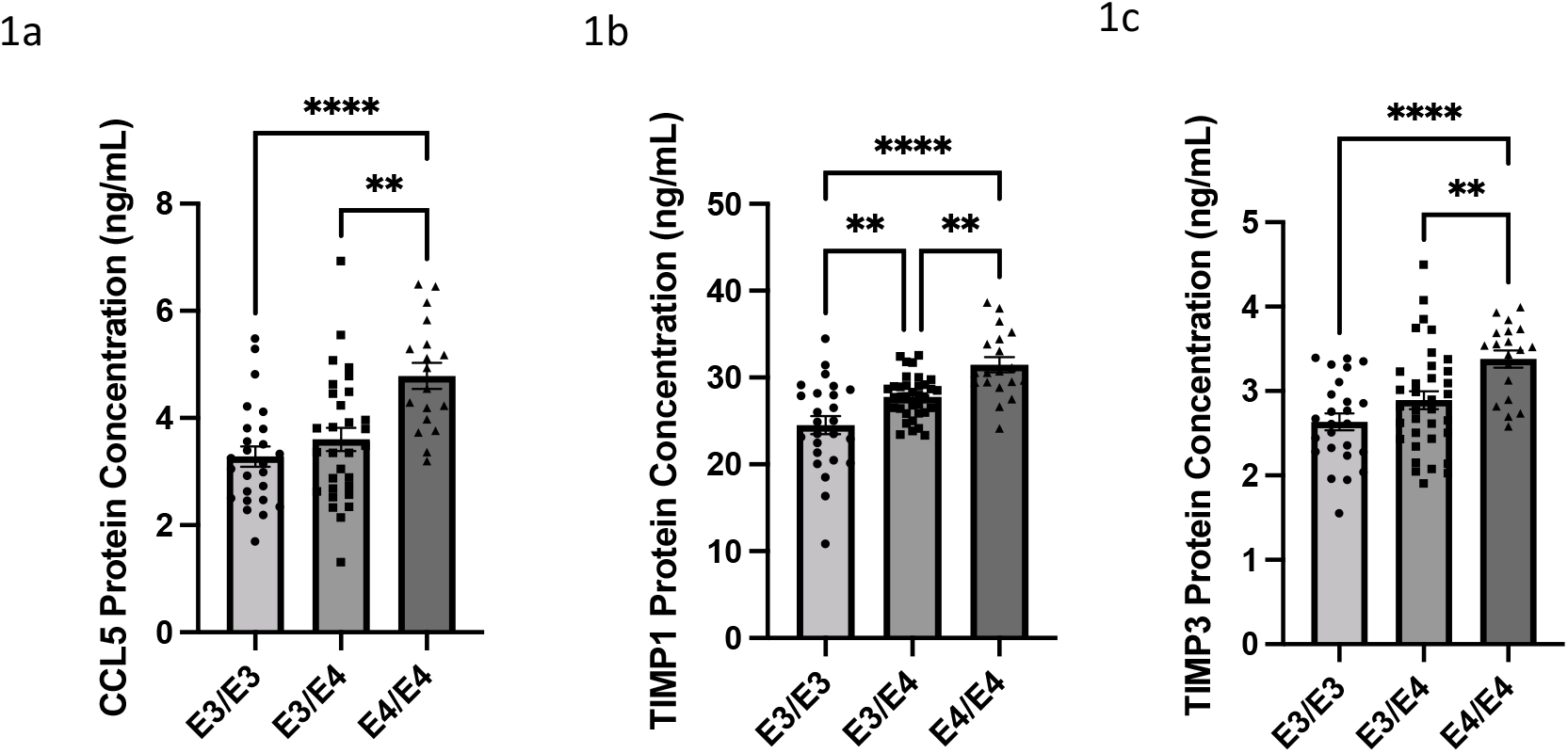
*APOE4/4* AD patients have significantly higher CCL5 protein concentrations detected in CSF as compared to *APOE3/3* and *APOE3/4* AD patients (*p* < .0001 and *p* = .0012, respectively, ANOVA with Tukey’s *post hoc* multiple comparisons), as measured by ELISA. *APOE4/4* AD patients also have elevated levels of TIMP-3 protein concentrations as compared to *APOE3/3* and *APOE3/4* AD patients (*p* < .0001 and *p* = .0068 respectively, ANOVA with Tukey’s *post hoc* multiple comparisons), as detected by ELISA. Both *APOE3/4* and *APOE4/4* AD patients have significantly higher TIMP-1 protein concentration, as compared to *APOE3/3* patients (*p* = .0049 and *p* = .0033, respectively, ANOVA with Tukey’s *post hoc* multiple comparison, n ≥ 18 CSF samples per genotype for all analytes).

**Figure 2.**
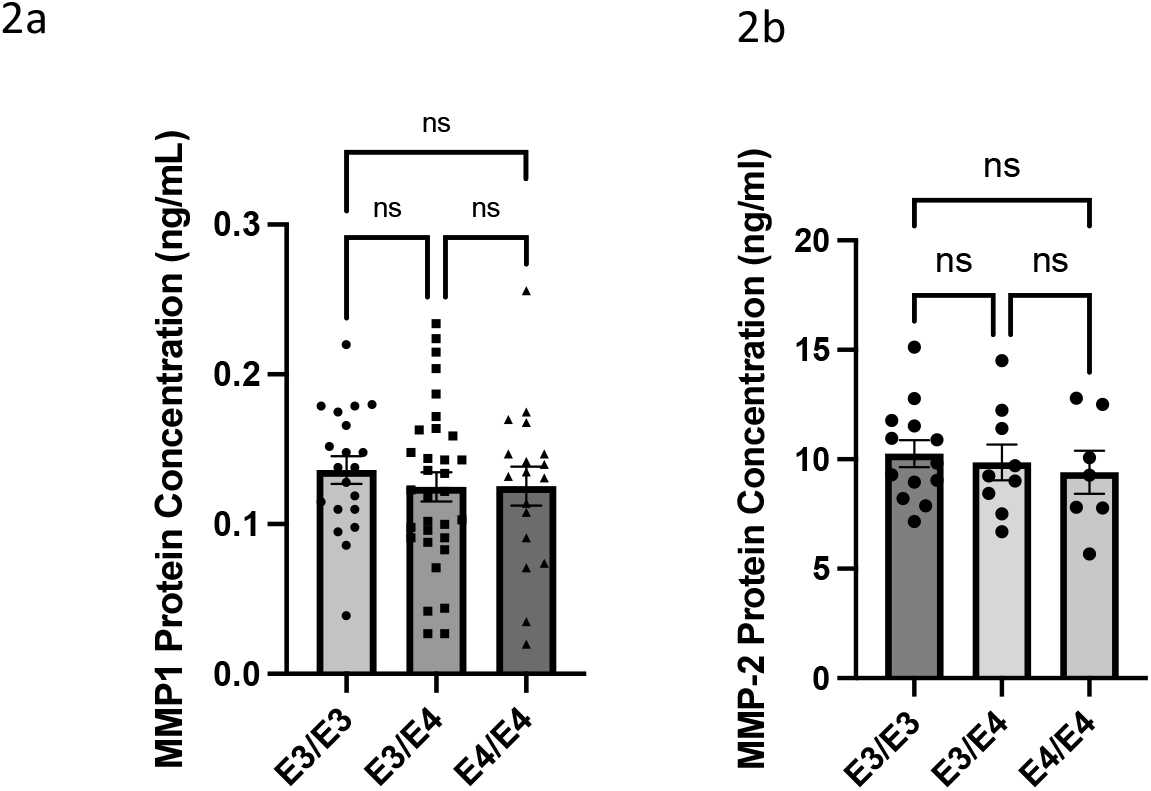
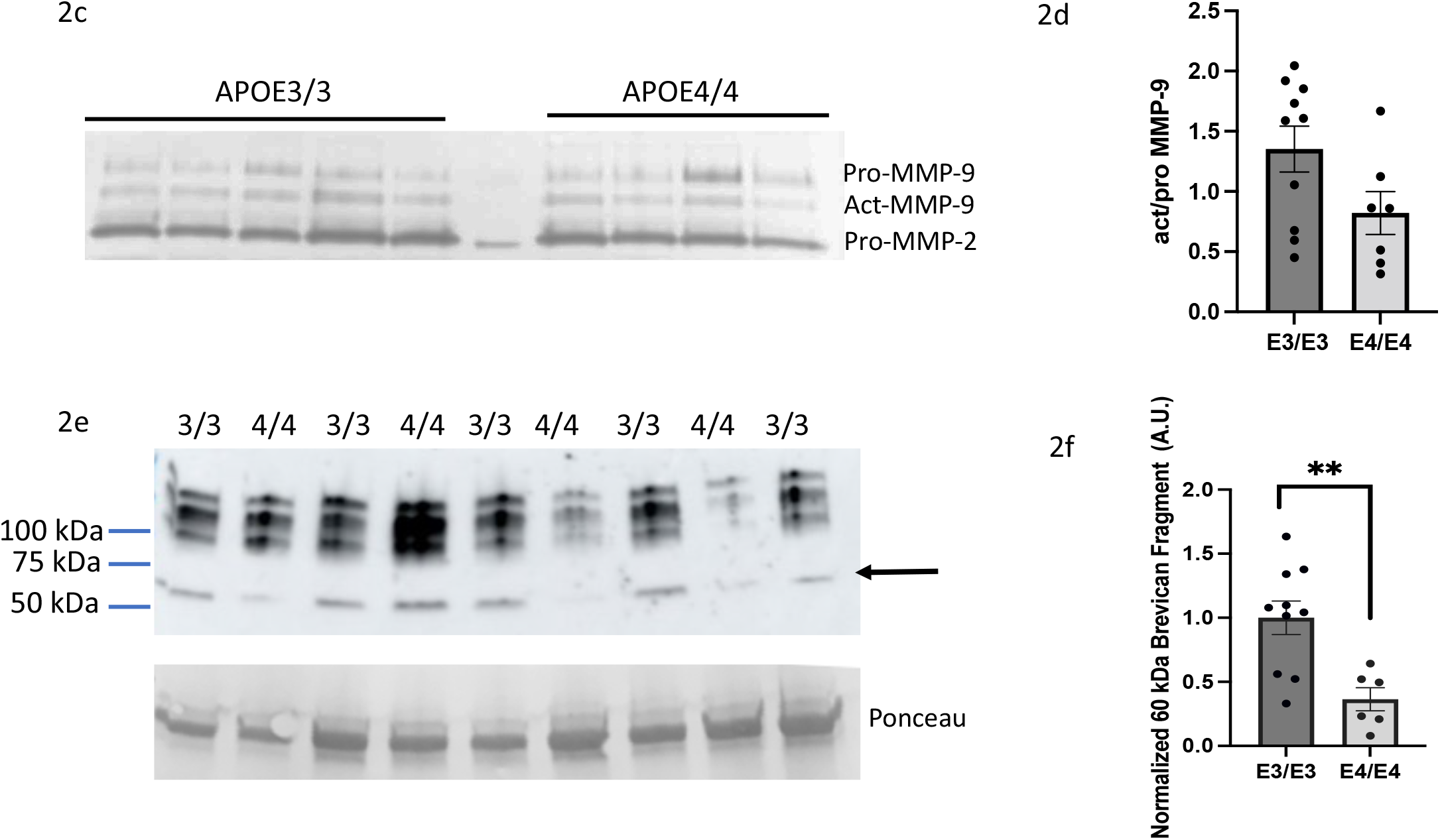
MMP-1 and MMP-2 protein levels (2a-b) do not show differences as a function of AD patient *APOE* genotype (n ≥ 18 CSF samples per genotype for MMP-1). Note fewer samples were analyzed for MMP2 because many pre-treatment samples had been depleted and thus 52-week post-placebo samples were instead used for this assay as described in the methods section. Representative gelatin substrate zymography and Western blot images on 10 post-placebo E3/E3 CSF samples and 7 post-placebo E4/E4 samples are also shown (2c and 2e). Densitometric data is shown in 2d and 2f respectively. The ratio of active to pro MMP-9 was non-significantly reduced in E4/E4 samples (*p*=0.07) but levels of a 50-60 kDa metalloproteinase brevican cleavage fragment (arrow) was significantly reduced in E4/E4 samples (*p*= 0.0039, Student’s *t* test). Note, one outlier was removed.

### Mice

Strains used for this study included C57BL/6J mice expressing human *APOE2, APOE3* or *APOE4* under the control of the endogenous murine *APOE* promoter, that have been previously validated (Xu *et al*., 1996; Sullivan *et al*., 1997). We also used strains from the Jackson Laboratory. Mice obtained from Jackson Laboratories included wild-type/C57BL6J background or CCR5 knockout (KO) mice on a C57BL6J background. *APOE4* TR mice were also crossed with wild-type or CCR5 KO mice to generate mice either with (1) a heterozygous TR of *APOE4* on a wild-type background or (2) a heterozygous TR of *APOE4* and a heterozygous KO of CCR5. Mice were housed four-five per cage. Food and water were provided *ad libitum*. Experiments were performed in accordance with National Institutes of Health guidelines and institutionally approved protocols (2016-1117 and 2018-0037). Cages were supplied with balconies and nesting materials for enrichment. Mice were anaesthetized with isoflurane and insensitivity to deep pain was confirmed prior to euthanasia by rapid decapitation.

### Primary astrocyte cultures

Primary astrocytes from 1-2 day old *APOE* TR mice were prepared as previously described (Lanfranco *et al*., 2021) and then stored in liquid nitrogen. For studies herein, 2 vials for each genotype were rapidly thawed and added to pre-warmed (37°C) medium (Gibco Minimal Essential Medium with 10% fetal calf serum and 1X penicillin streptomycin). Astrocytes were plated onto 6 well Costar plates. At confluence, the media was changed and supernatant samples were taken 12 hours subsequently for ELISAs.

### Brain lysates

Following euthanasia, hippocampi and cortices were micro-dissected. Regional lysates were prepared by lysis in immunoprecipitate buffer [50 mM Tris, pH 7.5, 150 mM NaCl, 0.1% sodium dodecyl sulfate, 1% octylphenoxypoly (ethyleneoxy) ethanol, branched, and 1X protease and phosphatase cocktail (Thermo Scientific 1861281)]. Lysates were sonicated for 10 seconds, placed on ice for 20 minutes, and centrifuged 15 minutes at 14,000 rpm at 4°C. Lysate supernatants were saved for protein analyses.

### ELISA and Western blot

Specific protein concentrations in CSF samples and tissue lysates were measured by ELISA (R & D Systems, Minneapolis MN), performed according to the manufacturer’s instructions, with the following changes. The incubation of sample with the antibody coated well was in the 4°C cold room overnight. In addition, the volume of sample added was 30 μl for murine lysates with 70 μl assay buffer. Note, that for each ELISA, all samples were run on a single plate to limit inter-experimental variability. For immunoblotting, 20 μl CSF was mixed with Laemmli sample buffer (Bio-Rad, Hercules, CA, USA, catalog #161-0737) containing 5% β-mercaptoethanol, and boiled for 5 minutes at 95 °C. Samples were subsequently separated by electrophoresis on precast gels (4-20% mini protein TGX gels, Bio-Rad catalog #456-1094) and transferred to nitrocellulose membranes (Trans-Blot Turbo Transfer, Bio-Rad, catalog #1704159). Membranes were probed with primary and secondary antibodies, and bands visualized by chemiluminescence as previously described (Alaiyed et al., 2019). We utilized a primary antibody to human brevican (1:1000, ThermoFischer PA5-4753). For densitometric normalization, since more than one gel was required to compare E3/E3 versus E4/E4 CSF samples by Western blot, we divided each band at the same molecular weight by the average of all E3/E3 band densities at that molecular weight for each individual image.

### Zymography

Gelatin (denatured collagen IV) substrate zymography was performed using precast gels from Invitrogen/Novux. 10% gels with 50 μl wells were loaded with 20 μl CSF and 20 μl zymogram sample buffer (BioRad). Samples were separated by electrophoresis and gels were then extracted and incubated for 30 minutes in renaturation buffer (Novux) followed by 3 days in development buffer (Novux) at 37°C. The relatively long development was necessary for the visualization of MMP-9 activity.

### T-Maze

Working memory was assessed in these mice using the T maze (Shoji *et al*., 2012). The task was performed by placing a mouse at the base of the T apparatus, and allowing the mouse to traverse down the length of the maze and explore either the left or right side of the T. Immediately following the mouse’s choice of goal arm, a door placed above each side of the maze was closed and the mouse was allowed to explore the side for 30s. The mouse was then removed before sanitizing the entire apparatus, after which the mouse was placed at the beginning of the T again to repeat the task. This version of the task does not involve utilizing a reward, and for each subsequent trial following the first, data is recorded as either a 1 or 0 to indicate whether the mouse spontaneously alternated arms of the maze or not. This task was repeated in random order for alternating cohorts of mice, and the data are presented as an average percent of alternation over the course of 4 trials.

### Fear Conditioning

Long-term hippocampal-dependent memory was assessed using fear conditioning (FC). Briefly, the training day for FC began with mice placed in the FC apparatus and allowed to explore the novel environment containing patterned walls for 180s. Following the acclimatization period, mice received a mild shock (0.5mA, 1s) at 180s, 240s, and 300s, before being removed from the apparatus after 360s. For the entirety of the training period, mice were video recorded and monitored using ANY-maze software, which measured bouts of freezing (defined as the mouse being immobile for >1s), the total duration of freezing, and latency to freeze. Long-term memory was assessed three days later when mice were placed in the same FC chamber with the contextual environment, and the same metrics were measured over a 180s period in the absence of floor shocks.

### In Vivo EEG Recordings

Telemetry recordings were performed with a DSI telemetry system connected to LabChart (AD Instruments) acquisition software via a Physiotel Bridge. We utilized mouse-sized implant transmitters which allowed for EEG and EMG recordings. The transmitter was placed subcutaneously in the lower back region and the electrodes were tunneled back towards the head. Surgery was performed by a D.V.M./Ph.D. veterinary surgeon (S.H. Hong) with sterile technique and general anesthesia continuous inhaled isoflurane), pre- and post-operative pain relief, and prophylactic antibiotics. Sutures were used to close the surgical wound and removed 10-14 days after surgery. A differential EEG configuration was recorded (L Frontal to R Parietal) from epidural screws, and EMG electrodes were placed in the cervical trapezius muscle. Video records were captured simultaneous with EEG. Signal was detected from implants (wirelessly) through a pad beneath the standard mouse housing units. Mice were able to see each other through clear housing units and are provided with enrichment. Signals were recorded and stored in LabChart format for offline data analysis. Data analysis included appropriate filtering and calculation of power in specific frequency ranges. After full recovery from surgery (3 days) low gamma power (20-55 Hz) was calculated for a 60-minute interval once every two hours during the dark/active phase (6 pm- 6 am and the average of power from n=6 one-hour recording epochs recorded for each day.

### Statistics

Sample size was based on power analyses for expected differences in human CSF and murine lysate values based on our prior published studies of inflammatory molecules for mu and sigma values (Conant *et al*., 1999; Alaiyed *et al*., 2020). For CSF analyses, the alpha value was 0.05 and desired power set at 0.80 as suggested at https://www.stat.ubc.ca/~rollin/stats/ssize/n2.html. All data were entered into a GraphPad Prism 8.0 program and statistical analysis was performed using Student’s unpaired *t*-test for two group comparisons or ANOVA, with post-hoc analyses as indicated, for comparisons of more than two groups. Significance was set at *p* < 0.05 and ROUT testing was performed to identify outliers.

## Results

### I. APOE4 is associated with elevated CSF levels of proteins that restrict neuroplasticity

To evaluate the possibility that CCL5 or other potential ECM effectors that may limit plasticity are elevated in association with *APOE4*, we first performed ELISA analysis of human CSF samples obtained from *APOE*-genotyped individuals with mild to moderate AD. Patient demographics are outlined in Table 1. As shown in Figure 1, *APOE4/4* AD patients have significantly higher CCL5 protein concentrations detected in CSF as compared to *APOE3/3* and *APOE3/4* AD patients (*p* < .0001 and *p* = .0012, respectively), as measured by ELISA. *APOE4/4* AD patients also have elevated levels of TIMP-3 protein concentrations as compared to *APOE3/3* and *APOE3/4* AD patients (*p* < .0001 and *p* = .0068, respectively), as detected by ELISA. Furthermore, both *APOE3/4* and *APOE4/4* AD patients have significantly higher TIMP-1 protein concentration, as compared to *APOE3/3* patients (*p* = .0049 and *p* = .0033, respectively). In our study sample, there was no correlation of CCL5, TIMP-1, or TIMP-3 with age (R^2^ = 0.00043, 0.0243, and 0.016 respectively)

**Table 1.** Baseline CSF sample patient demographics

| Genotype | Age (average +/- SEM) | Male/Female (n, %) |
| --- | --- | --- |
| *APOE3/3* | 72.24 +/- 1.48 | 14, 42.4% / 19, 57.6% |
| *APOE3/4* | 73.65 +/- 1.04 | 22, 43.1% / 29, 56.9% |
| *APOE4/4* | 69.87 +/-2.22 | 9, 39.1 % / 14,60.9 % |

### II. Specific matrix degrading molecules are not concomitantly increased in CSF samples from APOE4 positive individuals

Depending on the stimulus or disease, the expression and activity of matrix degrading molecules can change with TIMPs in a concordant or discordant fashion. To evaluate the possibility that MMPs might be concomitantly increased to abrogate effects of elevated TIMPs, we examined MMP levels in CSF samples from *APOE*-genotyped individuals. There was no difference in MMP-1 or MMP-2 protein levels as a function of AD *APOE* patient genotype in our sample group (Figures 2a and b). Note fewer samples were analyzed for MMP-2 because many pre-treatment samples had been depleted and thus 52-week post-placebo samples were instead used for this assay as described in the methods section. The post-placebo patient demographics are shown in Table 2. We also evaluated MMP-9 levels by ELISA (R & D Systems), and while we have previously detected MMP-9 in cortical brain lysates from humans with this ELISA (Alaiyed *et al*., 2020), in the CSF samples used for this study MMP-9 levels were below the detectable limits. We thus performed gelatin-based zymography for E4/E4 and E3/E3 individuals (Figure 2c). This assay allowed for quantitation of pro and active forms of MMP-2 and MMP-9. While we did not detect a significant difference in either, active MMP-9 levels appeared slightly diminished in E4/E4 individuals (Figure 2d, *p*= 0.07). Because gelatin-zymography is limited by a denaturation step that imparts activity to pro-forms and may also dissociate TIMPs from MMPs, we also looked at MMP-9 substrate cleavage as a function of APOE genotype. Results for brevican, which is cleaved by MMP-9 and other metalloproteases (Nakamura *et al*., 2000), are shown in Figures 2e and f. In brain extracts, particulate brevican may be membrane-linked and/or contribute to insoluble lattices including PNNs (Seidenbecher *et al*., 1995), while soluble brevican, the majority form, may also access the CSF compartment. Both forms may be cleaved. As shown in Figure 2e, our blots show high molecular weight soluble brevican as well as a previously described approximately 50-60 kDa metalloprotease generated cleavage fragment (Seidenbecher *et al*., 1995; Hussler *et al*., 2022). This proteolytic fragment is significantly increased in APOE3/3 as compared to APOE4/4 CSF samples (*p*= 0.0039, Student’s t test).

**Table 2.** 52-week post-placebo CSF sample patient demographics

| Genotype | Age (average +/- SEM) | Male/Female (n, %) |
| --- | --- | --- |
| *APOE3/3* | 75.5 +/- 2.4 | 5, 38.46% /8, 61.53% |
| *APOE3/4* | 75.4 +/- 1.97 | 5, 55.56% /4, 44.44% |
| *APOE4/4* | 70.85 +/- 6.04 | 4, 57.14% /3, 42.86% |

### III. Astrocytes from APOE4 TR mice secrete high levels of CCL5 and TIMP-1

To examine potential cellular sources of increased CCL5 and TIMP-1 expression as a function of *APOE* genotype, we also evaluated basal release of these molecules in astrocytes cultured from *APOE* TR mice. We focused on astrocytes since this cell type is numerous in the CNS and astrocytes can express both CCL5 and TIMP-1 (Crocker *et al*., 2006; Lanfranco *et al*., 2017). As shown by ELISA analyses of culture supernatants (Figure 3 a and b), astrocytes cultured from *APOE4/4* TR mice release significantly more CCL5 and TIMP-1 as compared to those from *APOE3/3* TR mice. The difference between CCL5 levels in E3/E3 and E4/E4 astrocytes is significant at *p*=0.0095 and the difference between E2/E2 and E4/E4 is significant at p< 0.0001 (ANOVA with Tukey’s multiple comparisons post hoc). For TIMP-1, in which we only examined E3/E3 and E4/E4 supernatants, the difference is significant at *p*< 0.0001 (Student’s t test).

**Figure 3.**
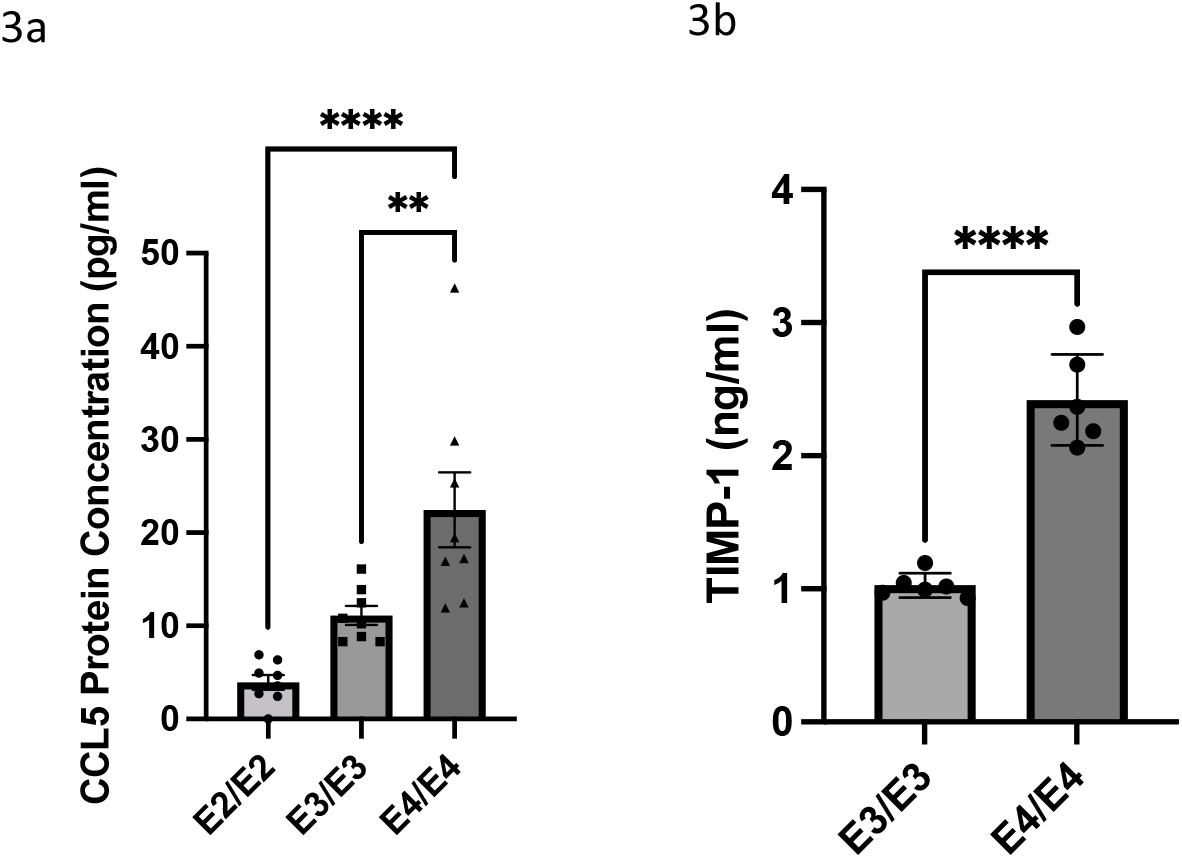
Astrocytes cultured from APOE3/3 TR and APOE4/4 TR mice show differential expression of CCL5 and TIMP-1. Shown are ELISA results of supernatants from APOE3/3 TR and APOE4/4 TR astrocytes. As shown in 3a and 3b respectively, CCL5 and TIMP-1 levels are increased in APOE4/4 TR murine astrocyte supernatants. CCL5 levels in E4/E4 are significantly increased as compared to E3/E3 or E2/E2 (p= 0.0001 and p<0.0095, ANOVA with Tukey’s *post hoc* multiple comparisons) and TIMP-1 levels were increased in E4/E4 as compared to E3/E3 (*p*< 0.0001, Student’s *t* test).

### IV. Gamma power is increased in CCR5 knockout mice

To evaluate the possibility that elevated CCL5 might influence gamma power in a manner observed in association with *APOE4* (Jones *et al*., 2019), we looked at gamma power in mice harboring a knockout of CCR5, the principal receptor for CCL5 in the brain. Our rationale for the use of this mouse was based on increased CCL5 expression with *APOE4* as well as prior work demonstrating increased pyramidal excitability in this model (Shen *et al*., 2022). As shown in Figure 4, gamma power (20-55 Hz) is significantly increased in CCR5 knockout mice as compared to wild type (*p* = .0063; n = 5 per genotype). Note: EEG activity was recorded over a 2-week period, and gamma power was calculated for a 60-minute interval once every two hours during the dark/active phase (6 pm- 6 am; average of power from n=6 one-hour recording epochs) and averaged per day (left panel). Days elapsed indicates EEG data recorded after animals had fully recovered from the implantation surgery and anaelgesia/anaesthesia (3 days).

**Figure 4.**
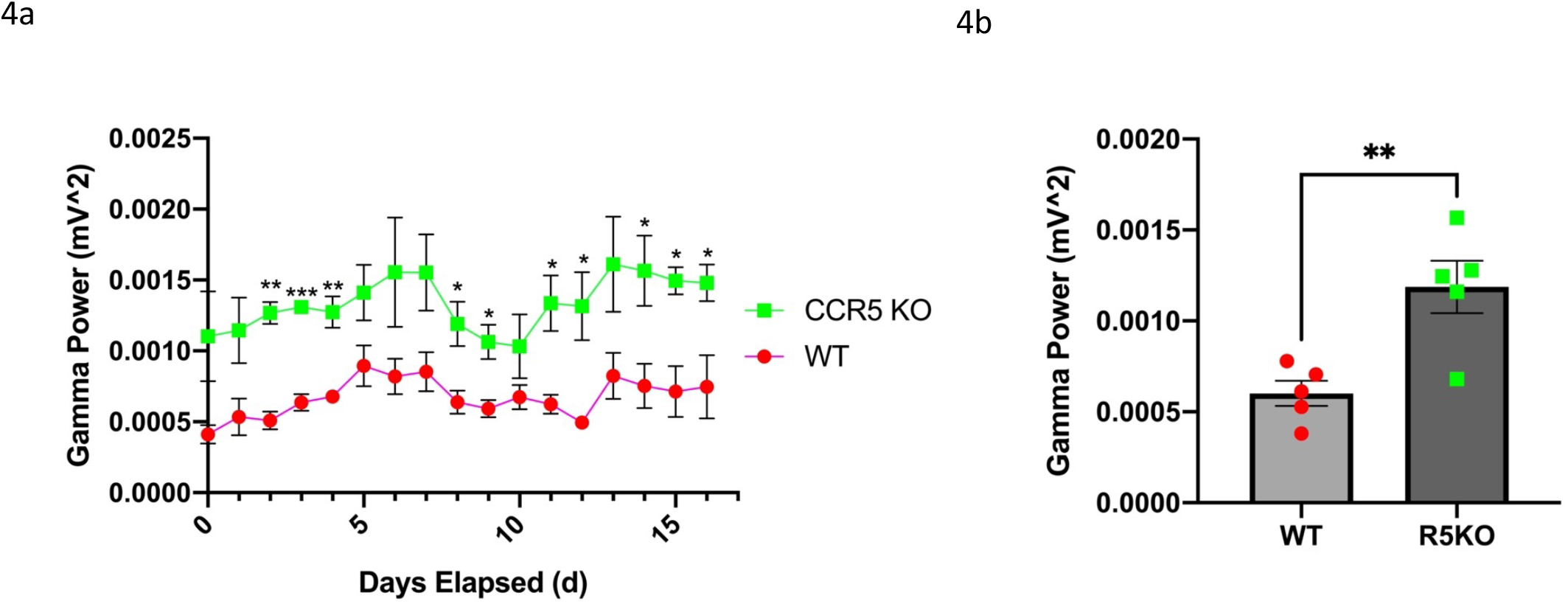
Gamma power (20 - 55Hz) is significantly increased in CCR5KO mice compared to age-matched wild-type mice (right panel; *p* = .0063; n = 5 per genotype). EEG activity was recorded over a 2-week period, and gamma power was calculated for a 60-minute interval once every two hours during the dark/active phase (6 pm- 6 am; average of power from n=6 one-hour recording epochs) and averaged per day (left panel). Days elapsed indicates EEG data recorded after animals had fully recovered from the implantation surgery.

### V. Gamma power (20 - 55Hz) is significantly increased in APOE4/CCR5KO heterozygous mice as compared to age-matched APOE4/wild-type mice

To determine whether CCR5 knockout could increase gamma power in the background of *APOE4* expression, we compared EEG activity in *APOE4* TR/ CCR5 KO heterozygotes to that in *APOE4* TR heterozygotes on a wild type background. As shown in Figure 5, gamma power (20 - 55Hz) is significantly increased in *APOE4*/CCR5KO heterozygous mice as compared to age-matched APOE4/wild-type mice (right panel; *p* = .0062; n = 3 per genotype). Gamma power was again calculated for a 60-minute interval once every two hours during the dark/active phase (6 pm- 6 am; average of power from n=6 one-hour recording epochs) and averaged per day (left panel). Days elapsed indicates EEG data recorded after animals had fully recovered from the implantation surgery.

**Figure 5.**
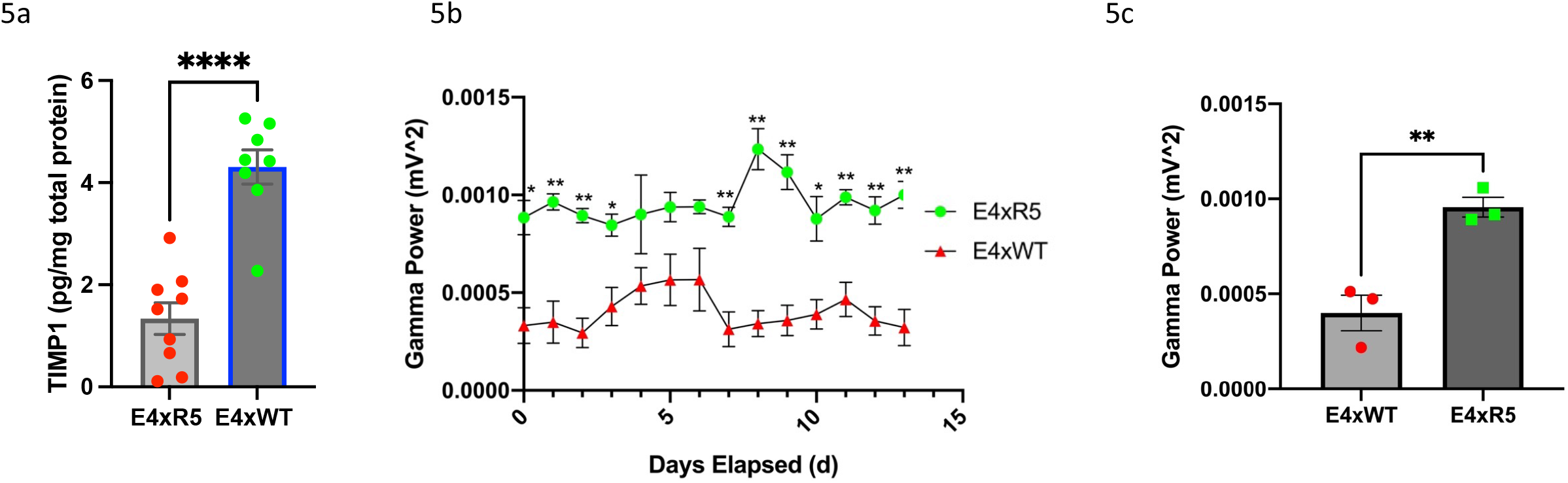
Gamma power (20 - 55Hz) is significantly increased in APOE4/CCR5KO heterozygous mice as compared to age-matched APOE4/wild-type mice (right panel; *p* = .0062; n = 3 per genotype). Gamma power was again calculated for a 60-minute interval once every two hours during the dark/active phase (6 pm- 6 am; average of power from n=6 one-hour recording epochs) and averaged per day (left panel). Days elapsed indicates EEG data recorded after animals had fully recovered from the implantation surgery.

### VI. APOE4/CCR5KO heterozygous mice show TIMP-1 normalization, as well as improved working memory and long-term memory in the T-maze and fear conditioning tasks

CCR5 is expressed on astrocytes (Lanfranco *et al*., 2017) and its engagement by endogenous ligands including CCL4 and CCL5 could contribute to their activation and expression of TIMP-1 (Crocker *et al*., 2006; Passos *et al*., 2009). We thus examined TIMP-1 levels in *APOE4* TR/ CCR5 KO heterozygotes and *APOE4* TR heterozygotes on a wild type background. As shown in Figure 6, TIMP-1 levels are reduced in hippocampal lysates from *APOE4* TR/ CCR5 KO heterozygotes as compared to APOE4 TR mice with normal CCR5 (6a). In addition, working memory and long-term memory performance, in which we saw significant increases in alternation on the T maze and freezing episodes during the recall portion of fear conditioning (Figure 6b and c), are consistent with improved memory in *APOE4* mice with reduced CCR5 expression.

**Figure 6.**
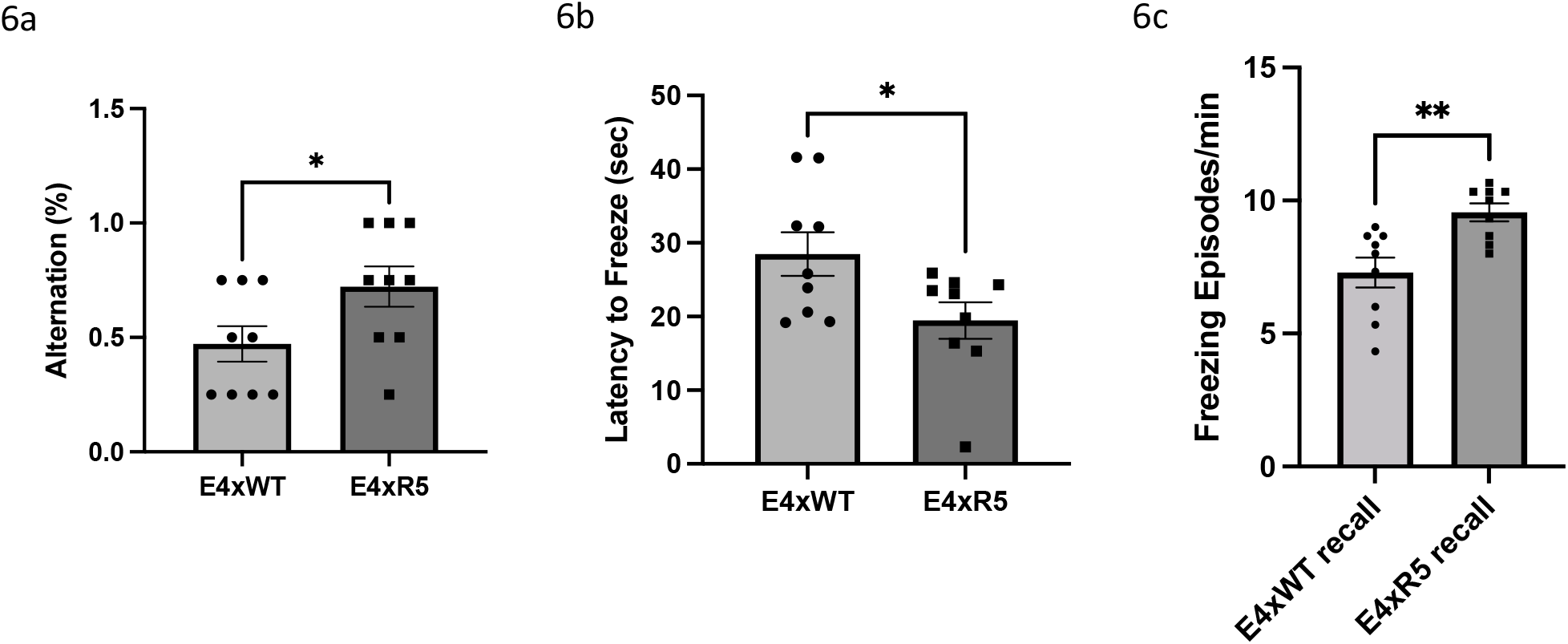
*APOE4*/CCR5KO heterozygous mice have reduced hippocampal TIMP-1 levels as well as improved working and long-term memory. Shown in Figure 6a are results from a TIMP-1 ELISA of hippocampal lysates from *APOE4*/CCR5 KO and APOE4/wild type heterozygotes. TIMP-1 levels are substantially reduced in the CCR5 heterozygous knockout lysates (*p*= 0.0001, Student’s *t* test) In Figures 6b and c, are results from the T-maze and fear conditioning tasks. APOE4/CCR5KO heterozygous mice have significantly increased alternation rate between the arms of the T-maze as compared to age-matched *APOE4*/WT heterozygous mice (*p* = .0485). *APOE4*/CCR5KO heterozygous mice had significantly more freezing episodes per minute in the recall trial of the fear conditioning task (*p* = .0032), and a significantly reduced latency to freeze on the trial day (p = .0328). No differences were observed in freezing episodes per minute during training (not shown).

## Discussion

In the present study, we found that APOE4 is associated with increased CCL5 and TIMP-1 levels in humans and TR mice, two molecules have been shown to negatively regulate neuroplasticity (Okulski *et al*., 2007; Zhou *et al*., 2016). Consistent with this, decreasing the CCL5 receptor CCR5 increases MAPK/CREB signaling, LTP and the temporal window for memory linking (Zhou *et al*., 2016; Shen *et al*., 2022). Endogenous CCL5 also blocks neuronal calcium oscillations (Meucci *et al*., 1998). In addition, TIMP-1 has been shown to inhibit MMP-9 dependent LTP in prefrontal cortex (Okulski *et al*., 2007).

CCL5 and TIMP-1 might also increase ECM deposition to less directly restrict plasticity. Indeed, CCL5 is linked to increased ECM deposition in liver and kidney and TIMP-1 inhibits the activity of ADAMs and MMPs that cleave constituents of both dense and diffuse ECM (Knight *et al*., 2019; Passman *et al*., 2021; Bonnard *et al*., 2022). We also show here that *APOE4* was associated with increased levels of TIMP-3. Previous work has shown that TIMP-3, which inhibits ADAM -10 and ADAM-17, reduces α-secretase mediated cleavage of amyloid precursor protein (Hoe *et al*., 2007). TIMP-3 levels are also increased in cortical tissues from human AD brain (Hoe *et al*., 2007).

Importantly, we observed no commensurate increase in MMP-1 or MMP-2 in *APOE4* human CSF, suggesting that elevated levels of TIMP-1 and TIMP-3 likely reduce overall proteolysis by the MMPs with which they interact. This could have consequences not only on ECM regulation but, in the setting of potential amyloid formation, on amyloid levels. Similar to select ADAMs, MMP-9 acts as an α-secretase and is also one of the few proteases that can also degrade fibrillar amyloid (Yan *et al*., 2006; Yin *et al*., 2006). In a tau model of AD, reducing ECM levels improves cognition (Yang *et al*., 2015). Moreover, in amyloid depositing mice engineered to express high levels of MMP-9, amyloid deposition is also reduced (Fragkouli *et al*., 2014; Yang *et al*., 2015). Similarly in human CSF, MMP-2 which shares substrate similarity with MMP-9, is negatively correlated with amyloid deposition as assessed by Pittsburgh compound B labeling (Sasaki *et al*., 2021). In a related study MMP-9 levels are reduced in AD patients as compared to controls (Mroczko *et al*., 2014). And though we acknowledge that other reports show that MMP-9 may be elevated at the blood brain barrier or in the brain parenchyma with human AD and aggressive mouse models of the same (Halliday *et al*., 2016; Weekman & Wilcock, 2016), whether this enzyme can ameliorate or exacerbate disease pathology is likely a function of quantity as well as localization. For example, increased expression of MMP-9 by activated microglia or pericytes at the blood brain barrier could have detrimental effects, while neuronal-derived and localized MMP-activity may target preferentially target PNNs and synaptic adhesion molecules to enhance plasticity (Tian *et al*., 2007). We did observe evidence for cleaved forms of neuron-associated molecules that influence plasticity in human CSF samples, and we suggest further studies may be warranted to examine additional neuronal substrates as a function of *APOE* genotype (Conant *et al*., 2015; Martin-de-Saavedra *et al*., 2022).

We also observed an effect of *APOE* genotype in primary astrocyte cultures (Figure 3), suggesting that the regulation of ECM proteolysis could occur in part through astrocytic mechanisms. Astrocytes cultured from E4 TR mice had increased supernatant levels of CCL5 and TIMP-1, suggesting that this cell type could contribute to changes seen in human CSF and murine brain lysates. Of interest, all human *APOE* isoforms have been shown to attenuate inflammation (Yin *et al*., 2019) and given that *APOE4* levels are reduced in comparison to other isoforms (Riddell *et al*., 2008), glial activation may be more prominent in the background of this particular isoform.

The most promising findings, however, were the normalization of TIMP-1 levels, EEG gamma power, and working memory through partial knockdown of CCR5. *In vivo* and *ex vivo* gamma power is enhanced by ECM attenuation (Lensjo *et al*., 2017; Bozzelli *et al*., 2020) and is thought to be important to working memory and attention. Gamma entrainment through sensory stimulation also improves cognition is a murine model of AD (Iaccarino *et al*., 2016) and has more recently been shown to reduce PNN levels (Venturino *et al*., 2021). Moreover, gamma power is reduced in APOE4 targeted replacement mice and the magnitude of reduction correlates with subsequent cognitive impairment (Jones *et al*., 2019). And though we did not directly compare wild type and APOE4 in a single surgery day or experimental cohort, as compared to wild type mice, the wild type APOE4 heterozygous animals had reduced gamma power (compare figures 6 and 7 in which wild type mice were generally over 0.005 mV^2^and E4 heterozygous mice were generally below this threshold). While we did not investigate the effects of CCR5 reduction in APOE3 TR mice, and we acknowledge that AD relevant effects may not be limited to *APOE4*, we note that it may be especially important to normalize gamma power in *APOE4* individuals.

Though our study is mechanistic in terms of demonstrating that a reduction in CCR5 normalizes *APOE4* relevant endpoints, a limitation is that we do not identify the downstream molecular effectors of CCR5-dependent normalization. Increased TIMP levels in E4 mice with normalization by knockdown of CCR5, and prior publications linking TIMPs to ECM deposition and reduced plasticity, support a potential role for altered ECM regulation in E4 mice as a contributor to behavioral and neurophysiological endpoints. This is supported by recently published work which shows chemokine and matrisome (ECM protein and associated factors) increases in human IPSC-derived *APOE4* astrocytes. This work also shows that *APOE4* microglia are enriched for ECM, chemokine and cytokine signaling pathways in multiple brain regions (Tcw *et al*., 2022). Additional mechanisms, however, may also be at play. For example, the effect of CCR5 antagonists on LTP are associated with an increase in MAPk/CREB activity in glutamatergic neurons that could also influence our endpoints (Zhou *et al*., 2016). Indeed, work from the Silva group has shown that pyramidal cell excitability is increased in the setting of CCR5 reductions and increased MAPk/CREB activity (Zhou *et al*., 2016). And as is the case for a reduction in inhibitory neuronal function, an increase in excitability of pyramidal neurons can enhance gamma oscillation power (Klemz *et al*., 2021). Future studies are thus warranted to assess the downstream mechanisms by which CCR5 normalizes gamma power and cognitive endpoints in the *APOE4* TR mice. Potential approaches include interventions that specifically target the ECM (Dubisova *et al*., 2022), and examination of ECM proteins in fixed brain tissue from individuals with and without *APOE4* with care to account for potential differences in glycan sulfation (Logsdon *et al*., 2022; Scarlett *et al*., 2022). In addition, since *APOE4* may increase the risk of recurrent episodes of major depression, a disorder often associated with reduced gamma power and cognitive deficits (Khalid *et al*., 2016; Papp *et al*., 2018; Alaiyed *et al*., 2019), future studies may be warranted to test CCR5 antagonists for this disorder.

In summary, we have demonstrated that CCL5 is increased in *APOE4* human CSF and brain lysates from *APOE4* TR mice and that biochemical, neurophysiological and behavioral deficits in heterozygous *APOE4* TR mice are normalized by heterozygous knockout of CCR5. We propose that it may be valid to consider use of the safe and well-tolerated CCR5 antagonist Maraviroc, which promotes plasticity and/or is neuroprotective in mouse models of HIV and stroke (Joy *et al*., 2019; Bhargavan *et al*., 2021), in select *APOE4* positive individuals.

## Acknowledgements

We would like to acknowledge outstanding veterinary support from the Department of Comparative Medicine and Dr. Patricia Foley. In addition, we would also like to apologize to investigators whose excellent work was not directly cited. We would also like to thank Tahiyana Khan and Dr. Patrick Forcelli for assistance in setting up the EEG recording software, Dr. Mark Burns for the use of his surgical area, and Christi Ann S Ng for shared resources. Katherine Conant, William Rebeck, and Griffin Greco received funds for support and/or supplies from NIH as follows: R01AG077002 (KC), R01 NS 100704 (GWR), and T32 NS 121780 (GAG).

CSF and plasma samples were collected with informed consent as a part of the Resveratrol clinical trial under Food and Drug Administration IND 104205, and registered at ClinicalTrials.gov (NCT01504854). All CSF and plasma samples were handled with strict anonymity throughout the study.

APOE genotype and demographic data used in the preparation of this manuscript were obtained from the University of California, San Diego Alzheimer’s Disease Cooperative Study (ADCS) Legacy database, supported by NIA U01AG010483 for the ADCS, ADCS funding sources, and funding by the ADCS. ADCS personnel including Dr. Robert Rissman contributed to the design and implementation of the ADCS and/or provided data but did not participate in analysis or writing of this report.

